# Male accessory gland size depends on genotypes but not larval density conditions in nutrition-rich environments

**DOI:** 10.1101/2025.05.21.655081

**Authors:** Bahar Patlar, Laura Japke, Sanja Moehring, Claudia Fricke

**Author notes:** Corresponds: CF; BP.

## Abstract

Accessory glands (AGs), responsible for producing seminal fluid, are critical for male reproductive success, especially under sperm competition. In *Drosophila melanogaster*, AG size is often used as a proxy for reproductive investment, but its variation in response to developmental environments, as well as distinct genotypes remain unclear. Here, we investigated how larval density without nutritional limitation per individual, genotype, and genotype-by-environment interactions (GEIs) influence AG size using 27 isogenic lines from the *Drosophila* Genetic Reference Panel (DGRP). While larval density significantly affected body size, with higher density producing smaller males even though their nutritional source is rich, we could not detect any effect on AG size, either in absolute terms or relative to body size. In contrast, AG size varied significantly among genotypes, with only minor GEI effects. Our findings suggest that AG size might be genetically determined and relatively insensitive to larval density cues under nutrient-rich conditions. These results highlight the limitations of using AG size as a sole proxy for reproductive investment and the importance of genotype in shaping male reproductive traits.

## Introduction

Post-mating sexual selection processes, particularly sperm competition [1], play a significant role shaping male reproductive success. In insects, accessory glands (AGs) within the male reproductive system produce the majority of seminal fluid proteins (SFPs) that mediate various post-mating functions, including sperm storage and release, suppression of female receptivity to subsequent matings, and enhanced paternity success in sperm competition [2–4]. As a result, AG size is often considered as a proxy for male reproductive investment, under the expectation that greater production of SFPs via larger AGs may enhance fertilization success. Empirical support for this comes from experimental evolution in *Drosophila melanogaster*, showing that selection for larger AGs increases some SFP production having role in sperm competition and improves male competitive fertility [5].

On the other hand, some indirect evidence also support the expectations. For example, AG depletion rates have been shown to evolve under different mating regimes in *D. melanogaster*, with males facing higher sperm competition depleting their AGs faster suggesting increased investment per mating [6]. Moreover, AG size is positively associated with mating frequency, duration and sperm displacement ability, implying a functional relevance with SFP amount that can depend on AG size [7]. Comparative studies also support the relationship between sperm competition intensity and AG size across species. Studies on some mammals and insects have demonstrated that species experiencing higher sperm competition tend to have larger AGs [8–10]. However, there are studies report contrasting results, possibly due to trade-offs between sperm and seminal fluid production in different mating systems [11].

Experimental evolution studies altering sex ratios to manipulate sperm competition intensity across generations has yielded mixed results on the relationships between sperm competition and AG size. In *D. melanogaster*, those studies found no significant changes in AG size under varying sex ratios across generations [6,12,13]. Similar findings were also observed in male seed beetles, *Callosobruchus maculatus* [14]. In contrast, in *Drosophila pseudoobscura*, populations exposed to higher competitive environments evolved larger AGs, correlating with increased mating frequency and offspring production [15], highlighting species-specific responses potentially shaped by differing SFP profiles and sperm morphologies [16,17].

Phenotypic plasticity, the ability of a genotype to produce different phenotypes in response to environmental conditions, is a critical mechanism enabling adaptation to varying selective pressures [18]. AG size shows plasticity in response to environmental conditions, including nutrition [19] and social cues of sperm competition during adult and developmental stages in different species including *D. melanogaster* [20–22].

Larval density, a critical environmental factor relevant in nature [23] affecting developmental trajectories, has been shown to influence life-history and sexual traits in adult *D. melanogaster* [24–26]. Males grown at low larval densities develop larger body size compared to those from high-density environments [27,28]. Larger males typically exhibit higher mating rates, increased remating success, higher production of key SFPs having role in sperm competition and enhanced offensive sperm competitiveness [27,29–31]. Thus, testis size and sperm production, as well as AG size are expected to respond plastically to the perceived sperm competition risk via larval crowding cues during development. So far studies have shown that while testis size shows no plastic response to larval density [21,32] conflicting results have emerged regarding AG size [21,22,32]. For example, Bretman et al. (2016) observed larger AGs in high-density conditions, even when accounting for body size. In contrast, Kapila et al. (2022) found smaller AGs under high density, both in absolute terms and relative to body size. Kapila et al. (2022) also found no response of larval density on AG size evolution across generations in their experimental evolution setup. While the results of those two studies relied on AG size in two-dimensions, this discrepancy is further complicated by Morimoto et al. (2022), who, using three-dimensional measurements, demonstrated a negative correlation between larval density and AG volume across a wider range of densities. Such methodological difference such as differences in larval density treatments, limiting nutritional resource in high densities [23], imaging techniques, or the genetic background of the studied populations could partly explain the discrepancies among studies. For example, a meta-analysis suggests that overall seminal fluid production tends to decline when nutrients are restricted, but responses vary across species [19].

A further layer of complexity is introduced by genotype-by-environment interactions (GEIs), which occur when genotypes differ in their response to environmental conditions. GEIs are particularly relevant for sexually selected traits, as they contribute to maintaining genetic variation under directional selection [33–35]. This variation, in turn, promotes the rapid evolution of reproductive traits such as SFPs and can even drive speciation [36–39].

In this study, we investigated how AG size in *D. melanogaster* responds to developmental larval density in the absence of nutritional limitation, and whether this response varies across genotypes. Using 27 isogenic lines derived from the *Drosophila* Genetic Reference Panel (DGRPs) [40], and rearing them under high and low larval density conditions with protein-rich food, we quantified both absolute and body size corrected AG size in two-dimensions.

This approach allowed us to disentangle the effects of genotype, GEIs, and larval density on variation in male reproductive investment during development.

## Material & Methods

### Study organism and maintaining conditions

The DGRP is a resource of isogenic lines derived from a single natural population, providing a powerful tool for exploring genotype-dependent trait variation [40]. We successfully examined 27 DGRP isolines (i.e., genotypes) for larval density (i.e., environment) effects on AG size. DGRPs were obtained from the Bloomington Stock Center (Bloomington, IN), and maintained on standard sugar-yeast-agar medium (SYA [41]) and at 12:12 light-dark cycle, 25 ±1 °C, and 60 ±10% humidity.

### Larval density manipulation and sampling

First instar larvae from each of the 27 DGRP lines were collected ∼24 hours after egg laying on grape-juice-agar plates (50 g agar, 600 ml red grape juice, 42.5 ml nipagin solution, 1.1 L water) overlaid with a thin layer of bakers’ yeast paste. To obtain larvae, hundreds of young adult flies from stock vials were placed in mating chambers containing grape-juice-agar plates in the morning, allowed to mate and oviposit for 24 hours, and then removed the next morning. The agar plates were inspected the following day, and first instar larvae were carefully collected. Larvae were distributed at random into one of two larval density conditions to develop: low density (50 larvae per vial) and high density (200 larvae per vial) treatment groups. To control for the effects of food resource limitation on overall body size, all larval density vials were prepared with *ad libitum* food enriched for protein (7 ml of SYA food including 150% yeast in 25-95 mm standard vials) where we reared the larvae until adult eclosion. Newly emerged virgin males were collected and flash-frozen at four days old when their AGs are mature [42]. Subsequently, up to 10 randomly selected males per larval density and genotype combination were dissected for AGs and wings.

### Measurements

Accessory glands were dissected in phosphate-buffered saline (PBS) by removing male genitalia and carefully cleaning away extraneous tissue. AGs were then mounted on microscope slides in ca. 15 µl PBS. To ensure consistent imaging, three layers of adhesive tape were put around the sides of the slides and AGs put in the middle. That allowed us to maintain AG arm alignment and prevent compression between the slide and coverslip. Images were captured using a microscope and camera (Zeiss Observer.Z1 with Axio Vision software release 4.8.2, Zeiss Microscopy) at 50× magnification. Concurrently, one wing per sample was mounted between a slide and coverslip and photographed using the same equipment.

As a proxy of AG size in two-dimension, AG area was measured by tracing one AG arm, applying a scaling factor, and exporting the area in square millimetres using TPS software [43]. Wing size as a proxy for body size was quantified as centroid size [44], calculated from the average of 11 landmark measurements in millimetres [45].

### Statistical analysis

All statistical tests were done using R (version 4.2.2) and RStudio (version 2024.12.0+467) [46,47]. R package *tidyplots* was used for data visualization [48]. Shapiro-Wilk tests were conducted to assess the normality of the wing and AG size measurements. Outliers in body size data were identified using *z*-scores (*z* > 3 or *z* <− 3) and AG size data was log-transformed to achieve normality. To analyse absolute AG size variation, we used the complete dataset of AG dissections that includes 7–10 males per genotype and larval density. Relative AG size, adjusted for body size, was analysed using a subset of individuals (3–10) for which both AG and wing measurements were available from the same individual.

First, linear mixed-effects models (LMMs) were fitted using the *lme4* package in R [49] to analyse body size and absolute AG size separately where larval density is the fixed factor, genotype (DGRP lines) and genotype-by-environment interactions (GEIs; DGRP lines nested within environment) are the random effects. These models were fitted using restricted maximum likelihood with a Gaussian family. Degrees of freedom for fixed effects were approximated using the Satterthwaite method implemented in the *lmerTest* package [50], which accounts for the uncertainty introduced by random effects and provides more accurate inference in mixed models.

Second, to analyse AG size relative to body size, we first conducted a simple Linear Model (LM) to assess the relationship between AG size as a response variable and body size as an explanatory variable at individual and genotype level. Considering the significant LM outcome, we then used a model comparison approach with LMMs including body size as a covariate and larval density as fixed effects, genotype and GEI as random effects. Model comparison was performed using AIC and BIC values and best fitted model outcome are reported using *lmerTest*.

Third, to estimate the effects of larval density on AG size while accounting for population structure, we fitted a Bayesian Mixed-Effects model using the *MCMCglmm* package in R. [51]. In the model, we similarly included larval density and body size as fixed effects. A random effect of genotype was included, with genetic relatedness among DGRP lines accounted for, which could otherwise obscure the true effects, via a correlation matrix constructed from a standardized genomic relationship derived from the DGRP resource (http://dgrp2.gnets.ncsu.edu; [40]). The model was run for 100,000 iterations, with a burn-in of 3,000, and a thinning interval of 5. Posterior distributions were summarized to obtain estimates and credibility intervals for fixed effects.

## Results

The Shapiro-Wilk test indicated a significant deviation from normality for both AG size (W=0.978, *p*=<0.001) and wing size (W=0.986, *p*=0.003). Log transformation resolved the normality issue for AG size (W=0.996, *p*=0.173). For centroid size, removal of two outliers out of 339 data points resulted in a normal distribution (W=0.992, *p*=0.061).

We first analysed body size and absolute AG size separately aiming to test effects of growing larvae under two density conditions for multiple genotypes, and their interactions (i.e., GEIs). The LMM including wing centroid size as the proxy for body size as the response variable showed a small but significant effect of larval density on body size that low larval density resulted slightly larger males (Table 1, Fig 1a). Random effect analyses showed high variance in body size among DGRP lines that explains around 70%, a negligible variance for GEI (0.02%) and a high variance within lines (i.e., residual variance that is 30%) (Table 1 & S1, Fig 1b). On the other hand, the LMM model for testing absolute AG size as the response variable showed no significant effect of larval density (Table 1, Fig 1c). Random effects for AG size also showed similar trends seen in body size variation, where they revealed low GEI variance, but higher variances between and within DGRP lines (Table 1 & S2, Fig 1d).

**Figure 1:**
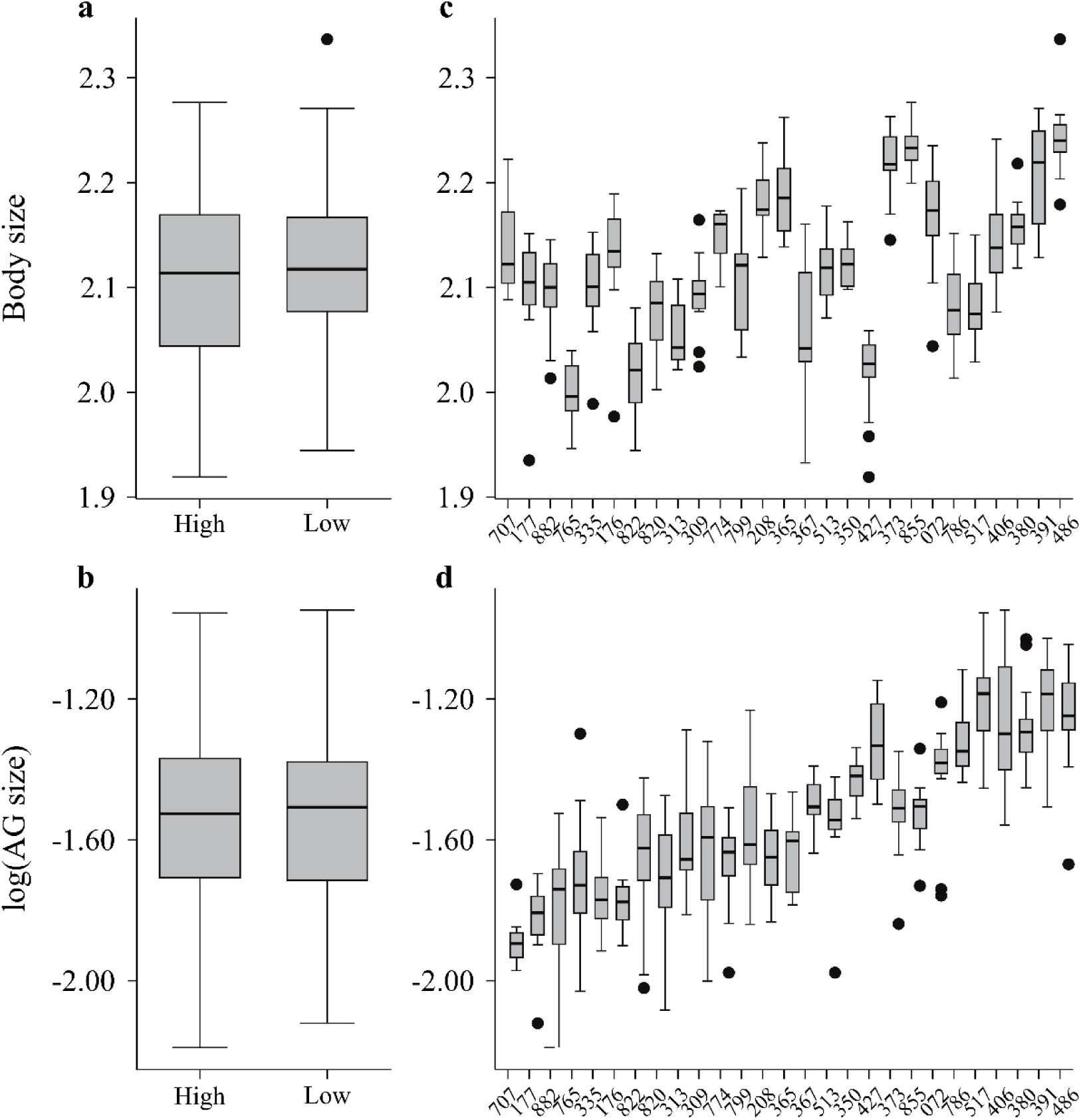
Distribution and mean values for body size (measured as wing centroid size) and absolute accessory gland (AG) size in relation to larval density and genotypes (i.e., *Drosophila* Genetic Reference Panel (DGRP) isolines). Left Panel includes boxplots show comparison of body size (a) and AG size (b) under low and high larval density conditions overall for all DGRPs. Right panel includes boxplots show comparisons of body (c) and AG size (d) values averaged across genotypes within environments, where genotypes are ordered by AG size in (d). DGRP lines reported as their original line (x-axis in c & d) identification numbers ([40]) Boxplots display the median and interquartile range.

**Table 1:** Effect estimates for body size (measured as wing centroid size) and accessory gland (AG) size in relation to larval density and genotypes (i.e., *Drosophila* Genetic Reference Panel (DGRP) isolines). The table provides outcome of linear mixed-effects model including larval density as a fixed factor. Random effects are included as variance components for genotype, genotype-by-environment interaction (GEI), and residuals, with the proportion of total variance was calculated for each random effect. For each trait, fixed-effect results include the estimate (Est), standard error (SE), degrees of freedom (DF, estimated using Satterthwaite’s approximation), *t*-value, and *p*-value. Statistical significance is indicated by *p*-values < 0.05 shown in bold.

| Effect type |  | Factors | Est. | SE | DF | *t*-value | *p*-value | % Var. |
| --- | --- | --- | --- | --- | --- | --- | --- | --- |
| Body size | Fixed | Larval density | 0.016 | 0.006 | 22.1 | 2.936 | **0.008** |  |
|  | Random | Genotype |  |  |  |  |  | 0.707 |
|  |  | GEI |  |  |  |  |  | 0.026 |
|  |  | Residual |  |  |  |  |  | 0.266 |
| AG size | Fixed | Larval density | 0.024 | 0.019 | 23.4 | 1.247 | 0.223 |  |
|  | Random | Genotype |  |  |  |  |  | 0.635 |
|  |  | GEI |  |  |  |  |  | 0.031 |
|  |  | Residual |  |  |  |  |  | 0.334 |

Previous studies have shown a positive correlation between body size and AG size in *D. melanogaster* [7,21]. This was also tested here using a LM including AG size as the response and body size as the explanatory variable. Results supported these previous studies showing that AG and body size are positively correlated at the individual level in each environment (low density: F1,163 =7.270, *p*=0.008, high density: F1,170 =4.633, *p*=0.033). When the same relationship was tested at the genotype-mean level, the pattern for positive relationship is stay clear (Fig 2), however no longer significant. Under low density conditions, the relationship between genotype means for AG and body size showed only a trend (F1,25 =3.52, *p*=0.072), while no association was detected under high density (F1,25 =1.02, *p*=0.323).

**Figure 2:**
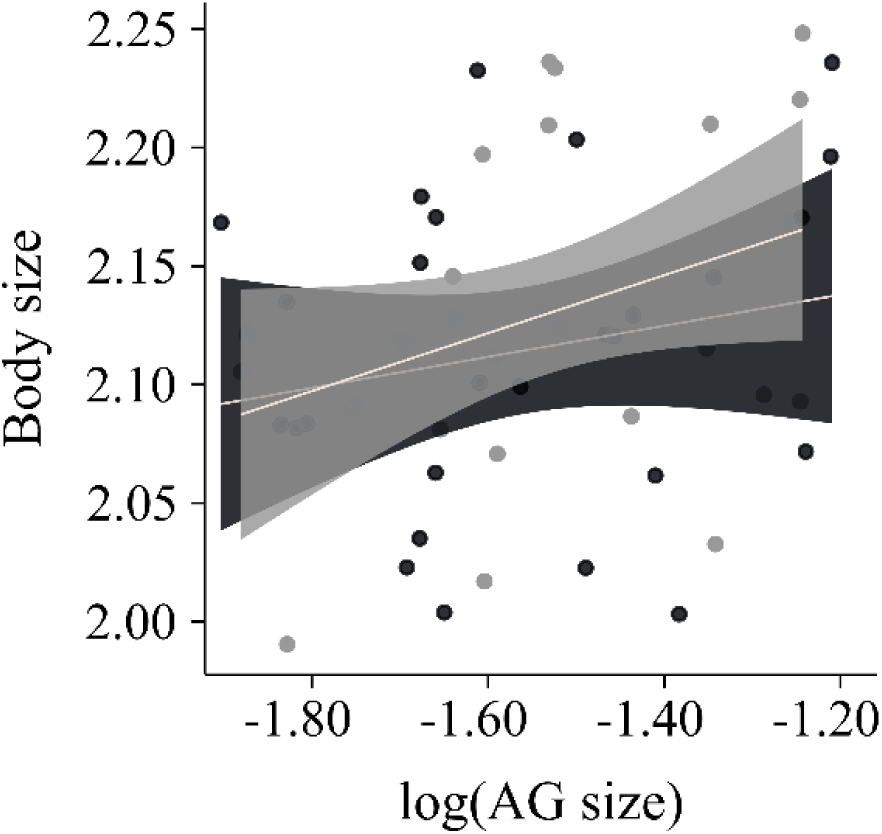
Scatterplot of genotypes (i.e., *Drosophila* Genetic Reference Panel (DGRP) lines) for average body size (measured as wing centroid size) and accessory gland (AG) size across larval densities. Scatterplot illustrating the relationship between body and AG size across high (black points) and low (grey points) larval density conditions for each genotype. Each point represents a DGRP line, with linear regression lines and 95% confidence intervals display for density conditions.

Considering individuals show a correlation at the individual level, to further explore the impact of larval density, genotype, and GEIs on AG size relative to body size we compared alternative models with varying combinations of fixed and random effects (Table S3). The full model with both body size and larval density as fixed effects, and genotype and GEIs as random effects had the lowest AIC and BIC values compared to reduced models, suggesting it provided the best balance between model complexity and goodness-of-fit (Table S3). The full model (Table 2) showed the effect of larval density remained statistically non-significant for AG size, suggesting no overall influence of larval density on AG size after also accounting for body size differences. The positive association between body size and AG size remained significant (*p*=0.013), confirming that this relationship holds at the individual level after accounting for genotypic and environmental sources of variation. The interaction effect is small while genotypes and within line variance show relatively higher variation for AG size that together explains 99% of total variance (Table 2).

**Table 2:** Effect estimates of body size (measured as wing centroid size), larval density, genotypes (i.e., *Drosophila* Genetic Reference Panel (DGRP) isolines) and genotype-by-environment interactions (GEI) on accessory gland (AG) size. The table reports fixed effects for body size and larval density, including the estimate (Est.), standard error (SE), degrees of freedom (DF), *t*-value, and *p*-value for AG size. Random effects include variance components for genotype, genotype-by-environment interaction (GEI), and residual variance, with the proportion of total variance was calculated for each random effect provided to indicate contribution to variability across components. Significant fixed effects (*p*-values < 0.05) are highlighted in bold.

| Effect Type | Effects | Est. | SE | DF | *t*-value | *p*-value | % Var. |
| --- | --- | --- | --- | --- | --- | --- | --- |
| Fixed | Body size | 0.478 | 0.191 | 329.597 | 2.501 | **0.0129** |  |
|  | Larval density | -0.010 | 0.019 | 24.207 | -0.527 | 0.6032 |  |
| Random | Genotype |  |  |  |  |  | 0.633 |
|  | GEI |  |  |  |  |  | 0.021 |
|  | Residual |  |  |  |  |  | 0.346 |

Finally, to account for genetic relatedness among DGRP lines, we fitted a *MCMCglmm* that showed a significant positive association between AG size and body size (posterior mean=0.484, 95% CI: 0.107–0.868, *pMCMC*=0.015) supporting the full model outcome generated with the LMM approach. Larval density had no significant effect on AG size (posterior mean=-0.016, 95% CI: -0.047–0.016, *pMCMC*=0.335). The Bayesian approach also showed that a substantial portion of the variation in AG size was attributable to differences among genotypes (G-structure) that was 0.0397 (95% CI: 0.019–0.065), while the residual variance (R-structure) was 0.0202 (95% CI: 0.0171–0.0236). This indicates that genetic background accounts for a meaningful proportion of the total phenotypic variance in AG size.

## Discussion

In this study, we investigated how larval density, genotype background, and genotype-by-environment interactions (GEIs) influence accessory gland (AG) size in *Drosophila melanogaster* by aiming to disentangle the effects of larval crowding from nutritional stress. Our results show that larval density affects body size slightly but significantly even in the absence of food limitation, with flies reared at low larval densities (50 larvae/vial) growing relatively larger than those from high larval densities (200 larvae/vial). However, we found no significant effect of larval density on AG size, either in absolute terms or relative to body size. Instead, AG size variation was found to be quite high among genotypes with no variation in GEIs.

Several previous studies have demonstrated that larval density with food restriction results in body size differences in *D. melanogaster* [25,27,28,52]. In our study, we aimed to minimize variation in body size across density conditions by providing *ad libitum* resources with high protein content and keeping slightly higher number of individuals in our low-density treatments compared to these studies. Despite these efforts, we still observed small but significant differences in average body size between low- and high-density treatments, highlighting larval density as a strong developmental factor shaping body size.

Consistent with earlier findings [7,21], we observed a significant positive correlation between body and AG size among individuals. Larger males tended to have larger AGs, suggesting that body size may act as a reliable proxy for reproductive trait investment in *D. melanogaster*. Larger body size due to low larval density has been associated with plastically increasing male harm to female [52] and female remating interval [53], phenotypes known to be mediated by SFPs produced by AGs, such as the Sex Peptide (SP) [54–56]. Thus, one can expect that larger males with larger AGs may produce and transfer higher amounts of such SFPs. However, contrary to predictions based on previous studies [21,22,32], AG size itself does not consistently scale with SFP production or transfer based on larval density treatments. For instance, Wigby et al. (2016) demonstrated that males reared at low larval density produce greater quantities of SP but transfer a smaller proportion of their SP reserves in each mating compared to high-density reared (smaller) males. Similarly, a recent larger scaled study has reported that despite producing slightly more SFPs overall, large males transferred fewer individual SFPs to females compared to small males, who transferred a greater quantity, including 10 specific SFPs in highly significantly higher abundances [57]. These recent findings collectively suggest that while AG size correlates with body size, it may not reliably predict functional ejaculate allocation depending on larval density, which could be regulated by more nuanced physiological and behavioural mechanisms.

On the other hand, it is worth noting that our low-density treatment may still entailed sufficient crowding to elicit higher allocation strategies to AG size. For example, some studies in which reported plastic increase in AG size and volume in higher larval density conditions implemented higher fold-change between their low and high densities then ours [22,32]. Thus, these discrepancies in our findings may stem from differences in larval density treatment designs. On the other hand, as already mentioned those studies also restricted food resource in their high-density treatments that may result in strategic investment during growth of different reproductive tissues. Nutrient limitation causes a moderate but taxonomically widespread reduction in male ejaculate traits, with seminal fluid quantity being most strongly affected, followed by sperm quantity, while sperm quality traits like viability and morphology are less consistently impacted [19]. In addition, theories of sperm competition predict diminishing returns to increasing sperm competition risk or intensity [58,59], and empirical evidence supports this for sperm number and size in *D. melanogaster* [60]. This framework might explain the limited plasticity in AG size observed in our study, as even our low density conditions could have reached a threshold of diminishing returns for effective sperm competition intensity [23].

Alternatively, the cellular composition of AGs may itself be a target in our larval density settings. The diversity, number, size and arrangements of specialized cell types within the AG could respond to developmental conditions like larval density without altering overall gland size. The *D. melanogaster* AG is composed of specialized cell types (main cells and secondary cells) with distinct roles in SFP production [61]. Indeed, in the *Drosophila* genus, the number and arrangement of specialized AG cells is known to evolve rapidly [61]. To better understand the relationship between AG size, the cellular components and reproductive success depending on larval density, e.g. how female post-mating responses change needs further experiments.

Both the LMM and Bayesian models confirm that genetic background is a significant factor in determining variation in AG size among DGRP lines that are known to have distinct genotypes and phenotypes [40,62]. Substantial genetic variation has also been reported for body size and several other SFP related phenotypes such as gene expression, female remating rate and paternity success in DGRP lines [28,63,64]. These findings reinforce the longstanding paradox of why male sexually selected traits such as AG size, retain high genetic variation despite the expectation of strong directional selection. The persistence of variation in male sexually selected traits can be attributed to the existence of GEI due to their condition-dependent natures [34,65]. However, we found a negligible contribution of GEI to the total variation in AG size plasticity suggest that AG size is relatively robust among genotypes across the larval densities. This is also in line with previous work showing no GEI for gene expression in DGRPs for SP and other SFPs involving sperm competition [28]. Therefore, maintaining genetic variation for AG size may not be driven by GEIs where population density shows heterogeneity in nature, but by several alternative mechanisms may be involving.

One explanation is that sperm competition may not exert strong selective pressure at the population level, particularly in large populations, due to the non-transitivity (i.e., the outcome of a given male’s sperm competition depends on the specific rival male’s genotype) [66,67]. This rock–paper–scissors dynamic means that there is no single optimal genotype, which in turn weakens net selection pressure and allows multiple genotypes (and thus variation in traits like AG size) to persist in the population. Moreover, each time in sperm competition a small number of males compete against each other and represent a subset of the genetic variation present in a population that overall decreases net selection intensity [68,69]. In addition, males from populations evolved under high sperm competition risk did not develop larger glands, but they exhibited faster depletion of AG reserves (measured as shrinking in size) over successive matings than males from low competition populations [6,13] suggesting organ size itself may not be target.

Additional evidence points to alternative selective forces or sperm competition as a weak force shaping AG traits under some circumstances. For example, natural genetic variation in body size of *D. melanogaster* was found to be not correlated with variation in sperm competitive ability, implying that larger males (and by extension possibly larger glands) do not consistently outcompete smaller males in fertilization [70]. Finally, a comparative perspective from other taxa underscores that AG evolution can be driven by factors other than sperm competition. In ray-finned fishes, for instance, lineages with male parental care evolved glands much more rapidly than those without, whereas high sperm competition actually showed a trend of slower AG evolution [71].

In conclusion, AG size appears largely genetically determined and resistant to larval density conditions under a nutrient-rich environment, or it responds by adjusting other structures instead of gross size such as cell number and size. Thus, future studies should adopt more integrative approaches. First, our measurement of AG size was limited to two-dimensional estimates, however, advancements in imaging technology (e.g. micro-CT scanning or confocal microscopy) can capture the true three-dimensional volume and structure of the AGs that was successfully implemented recently in *D. melanogaster* [32]. Second, we focused on a single developmental stage and environmental condition; indeed, longitudinal studies tracking AG size across multiple life stages and environmental gradients would provide a more comprehensive understanding of its plasticity. Finally, AG size is a very crude measure, and more in-depth analyses are needed such as investigating the relationship between SFP gene expression and protein amount that is produced by AGs, e.g., using single cell RNA transcriptome techniques. Such integrative approaches are needed to clarify and better understand the links between AG morphology, SFP production, cellular function, and fitness that shape the male reproductive organ evolution and reproductive success.

## Acknowledgement

This research was [partly] funded by the German Research Foundation (DFG) as part of the CRC TRR 212 (NC³) – Project number 316099922. Claudia Fricke was supported by a DFG funded Heisenberg fellowship (FR 2973/5-1 and Fr 2973/11-1) while conducting this work.

