## Supplementary material for "Male accessory gland size depends on genotypes but not larval density conditions in nutrition-rich environments": Supp: AG_size_MS_Supp.pdf

CF https://orcid.org/0000-0002-0691-6779

BP https://orcid.org/0000-0002-0442-9061

**Table S1: Summary of body size data (measured as wing centroid size) for *Drosophila* Genetic Reference Panel (DGRP) lines grown under high and low larval density conditions.** For each DGRP ID, the number of individuals measured (N), the mean wing centroid size, and the standard deviation (SD) are reported.

| | | High larval density | | | Low larval density | | |
|---|---|---|---|---|---|---|---|
| | **DGRP ID** | **N** | **Wing size** | **SD** | **N** | **Wing size** | **SD** |
| **1** | 176 | 3 | 2.135 | 0.042 | 5 | 2.119 | 0.083 |
| **2** | 177 | 5 | 2.083 | 0.086 | 6 | 2.105 | 0.030 |
| **3** | 208 | 7 | 2.171 | 0.022 | 6 | 2.197 | 0.041 |
| **4** | 309 | 3 | 2.120 | 0.039 | 8 | 2.081 | 0.035 |
| **5** | 313 | 8 | 2.035 | 0.016 | 9 | 2.071 | 0.027 |
| **6** | 335 | 7 | 2.083 | 0.054 | 5 | 2.117 | 0.020 |
| **7** | 350 | 4 | 2.121 | 0.030 | 5 | 2.124 | 0.021 |
| **8** | 365 | 7 | 2.179 | 0.027 | 3 | 2.209 | 0.064 |
| **9** | 367 | 6 | 2.023 | 0.045 | 6 | 2.087 | 0.066 |
| **10** | 373 | 7 | 2.203 | 0.033 | 5 | 2.236 | 0.024 |
| **11** | 380 | 7 | 2.145 | 0.019 | 7 | 2.168 | 0.028 |
| **12** | 391 | 6 | 2.196 | 0.060 | 4 | 2.220 | 0.041 |
| **13** | 406 | 6 | 2.170 | 0.044 | 6 | 2.115 | 0.034 |
| **14** | 427 | 9 | 2.003 | 0.044 | 9 | 2.033 | 0.026 |
| **15** | 486 | 8 | 2.236 | 0.023 | 6 | 2.248 | 0.052 |
| **16** | 513 | 5 | 2.099 | 0.022 | 7 | 2.131 | 0.035 |
| **17** | 517 | 8 | 2.072 | 0.038 | 9 | 2.093 | 0.030 |
| **18** | 707 | 3 | 2.168 | 0.062 | 5 | 2.121 | 0.031 |
| **19** | 72 | 6 | 2.129 | 0.048 | 5 | 2.210 | 0.026 |
| **20** | 765 | 7 | 2.004 | 0.030 | 5 | 1.990 | 0.035 |
| **21** | 774 | 5 | 2.151 | 0.023 | 6 | 2.146 | 0.029 |
| **22** | 786 | 7 | 2.062 | 0.033 | 10 | 2.096 | 0.043 |
| **23** | 799 | 9 | 2.101 | 0.055 | 4 | 2.112 | 0.018 |
| **24** | 820 | 8 | 2.063 | 0.048 | 7 | 2.091 | 0.031 |
| **25** | 822 | 5 | 2.023 | 0.040 | 8 | 2.017 | 0.044 |
| **26** | 855 | 8 | 2.232 | 0.022 | 6 | 2.234 | 0.021 |
| **27** | 882 | 8 | 2.082 | 0.043 | 3 | 2.128 | 0.010 |

**Table S2: Summary of accessory gland (AG) size measurements for *Drosophila* Genetic Reference Panel (DGRP) lines grown under high and low larval density conditions.** For each DGRP ID, the number of individuals measured (N), the mean AG size, and the standard deviation (SD) are reported. Values outside parentheses represent calculations for all individuals, while values inside parentheses represent measurements from individuals for whom both wing and AG sizes were recorded.

| | | High larval density | | | Low larval density | | |
|---|---|---|---|---|---|---|---|
| | **DGRP ID** | **N** | **AG size** | **sd** | **N** | **AG size** | **sd** |
| **1** | 176 | 9(3) | -1.829(-1.814) | 0.119(0.081) | 10(5) | -1.699(-1.733) | 0.243(0.147) |
| **2** | 177 | 9(5) | -1.806(-1.766) | 0.059(0.047) | 10(6) | -1.88(-1.878) | 0.196(0.136) |
| **3** | 208 | 10(7) | -1.659(-1.67) | 0.169(0.132) | 9(6) | -1.606(-1.631) | 0.113(0.131) |
| **4** | 309 | 9(3) | -1.458(-1.464) | 0.186(0.148) | 10(8) | -1.654(-1.695) | 0.195(0.197) |
| **5** | 313 | 10(8) | -1.678(-1.654) | 0.174(0.169) | 10(9) | -1.589(-1.585) | 0.108(0.114) |
| **6** | 335 | 10(7) | -1.835(-1.813) | 0.148(0.102) | 8(5) | -1.693(-1.678) | 0.09(0.093) |
| **7** | 350 | 8(4) | -1.467(-1.416) | 0.139(0.071) | 9(5) | -1.519(-1.443) | 0.145(0.063) |
| **8** | 365 | 10(7) | -1.676(-1.687) | 0.081(0.087) | 8(3) | -1.531(-1.51) | 0.053(0.057) |
| **9** | 367 | 10(6) | -1.489(-1.524) | 0.109(0.02) | 10(6) | -1.437(-1.469) | 0.152(0.088) |
| **10** | 373 | 9(7) | -1.499(-1.490) | 0.09(0.101) | 10(5) | -1.53(-1.563) | 0.146(0.167) |
| **11** | 380 | 10(7) | -1.344(-1.3) | 0.166(0.139) | 9(7) | -1.261(-1.254) | 0.095(0.108) |
| **12** | 391 | 7(6) | -1.211(-1.206) | 0.148(0.161) | 8(4) | -1.246(-1.253) | 0.176(0.175) |
| **13** | 406 | 9(6) | -1.244(-1.225) | 0.13(0.129) | 9(6) | -1.352(-1.311) | 0.213(0.253) |
| **14** | 427 | 10(9) | -1.383(-1.306) | 0.271(0.127) | 9(9) | -1.342(-1.342) | 0.111(0.111) |
| **15** | 486 | 9(8) | -1.209(-1.223) | 0.108(0.107) | 10(6) | -1.243(-1.291) | 0.173(0.216) |
| **16** | 513 | 10(5) | -1.563(-1.583) | 0.205(0.229) | 10(7) | -1.535(-1.538) | 0.046(0.035) |
| **17** | 517 | 10(8) | -1.239(-1.187) | 0.171(0.148) | 10(9) | -1.246(-1.228) | 0.121(0.114) |
| **18** | 707 | 8(3) | -1.903(-1.856) | 0.147(0.123) | 7(5) | -1.874(-1.903) | 0.186(0.042) |
| **19** | 72 | 8(6) | -1.435(-1.502) | 0.211(0.196) | 7(5) | -1.348(-1.329) | 0.066(0.07) |
| **20** | 765 | 10(7) | -1.649(-1.615) | 0.172(0.188) | 10(5) | -1.828(-1.837) | 0.192(0.129) |
| **21** | 774 | 10(5) | -1.677(-1.679) | 0.158(0.175) | 10(6) | -1.64(-1.667) | 0.102(0.109) |
| **22** | 786 | 10(7) | -1.410(-1.356) | 0.241(0.079) | 10(10) | 1.287(-1.287) | 0.111(0.111) |
| **23** | 799 | 10(9) | -1.610(-1.615) | 0.127(0.133) | 9(4) | -1.594(-1.52) | 0.223(0.267) |
| **24** | 820 | 10(8) | -1.660(-1.630) | 0.12(0.109) | 9(7) | -1.753(-1.791) | 0.196(0.18) |
| **25** | 822 | 9(5) | -1.692(-1.684) | 0.14(0.192) | 10(8) | -1.604(-1.636) | 0.176(0.179) |
| **26** | 855 | 10(8) | -1.612(-1.523) | 0.197(0.046) | 10(6) | -1.524(-1.525) | 0.116(0.136) |
| **27** | 882 | 10(8) | -1.818(-1.834) | 0.183(0.203) | 10(3) | -1.637(-1.741) | 0.167(0.217) |

**Table S3: Summary of model comparisons assessing the effects of larval density (Environment), *Drosophila* Genetic Reference Panel (DGRP) isolines (Genotypes), their interactions (GEIs), and body size (measured as wing centroid size) on accessory gland (AG) size.** The table includes the model names with their fixed and random effects, Akaike Information Criterion (AIC), Bayesian Information Criterion (BIC), and key findings. The Null model serves as a baseline for comparison, with only genotype as a random effect.

| *Model Name* | *Fixed Effects* | *Random Effects* | *AIC* | *BIC* | *Key Findings* |
|---|---|---|---|---|---|
| *Null* | None (~ 0) | Genotype | -152 | -144 | Baseline model for comparison |
| *Simple* | Wing centroid size | Genotype | -260 | -245 | Body size significant ($p$=0.009) |
| *Relative* | Environment, Wing size | Genotype | -252 | -233 | Body size significant ($p$=0.007), Environment not significant |
| *Relative & interaction* | Environment, Wing size | Genotype, GEI | -251 | -228 | Body size significant ($p$=0.013), Environment not significant |
| *MCMCglmm* | Environment, Wing size | Genetic relatedness matrix | | | DIC = -319.96<br>Body size significant (*pMCMC*=0.013), Environment not significant |
