## Supplementary material for "Male accessory gland size depends on genotypes but not larval density conditions in nutrition-rich environments": Supp: Supplementary_script_ AG_Centroid_analysis.R

```
# Clear Environment and Load Libraries
rm(list = ls())
getwd()

# Load Required Libraries
library(readr)
library(dplyr)
library(tidyplots)
library(Rmisc)
library(lme4)
library(lmerTest)
library(car)
library(MCMCglmm)
library(Matrix)
library(phyr)
library(tidyr)
library(ggplot2)

#------------------------------------------------------------------------------
# 1. LOAD AND CLEAN DATA
#------------------------------------------------------------------------------

# Load Data
AG <- read.table("AG_area_23072024.txt", header = TRUE, sep = "")
Centroid <- read.table("Centroid_size_23072024.txt", header = TRUE, sep = "")

# Exclude LineID "DGRP_18" missing from Centroid data
AG <- AG %>% filter(LineID != "DGRP_18")

# Merge datasets
data <- merge(AG, Centroid, by.id = "ID", all.x = TRUE)

# Exclude lines with <3 Centroid observations
centroid_count <- aggregate(Centroid ~ Env + LineID, data = data, FUN = function(x) sum(!is.na(x)))
lines_to_exclude <- unique(centroid_count$LineID[centroid_count$Centroid < 3])
data_filtered <- data[!data$LineID %in% lines_to_exclude, ]

#------------------------------------------------------------------------------
# 2. DATA TRANSFORMATION AND OUTLIER DETECTION
#------------------------------------------------------------------------------

# Log-transform variables
data_filtered$logag <- log(data_filtered$AG_area)
data_filtered$logbs <- log(data_filtered$Centroid)

# Outlier detection for Centroid
data_filtered <- data_filtered %>%
  mutate(z_score = ifelse(is.na(Centroid), NA, (Centroid - mean(Centroid, na.rm = TRUE)) / sd(Centroid, na.rm = TRUE)))

outliers <- data_filtered %>% filter(abs(z_score) > 3)
data_filtered <- data_filtered %>% filter(!image_name %in% outliers$image_name)

# Final data for models (no missing Centroid)
data_final <- data_filtered %>% filter(!is.na(Centroid))

#------------------------------------------------------------------------------
# 3. SUMMARY STATISTICS
#------------------------------------------------------------------------------

# Descriptive stats
Descriptives_log_AG <- summarySE(data_filtered, "logag", c("Env", "OriginalID"), na.rm = TRUE)
Descriptives_abs_AG <- summarySE(data_filtered, "AG_area", c("Env", "OriginalID"), na.rm = TRUE)
Descriptives_BS     <- summarySE(data_filtered, "Centroid", c("Env", "OriginalID"), na.rm = TRUE)

# Merge for correlation plots
cordat <- merge(Descriptives_log_AG, Descriptives_BS, by = c("Env", "OriginalID"))

#------------------------------------------------------------------------------
# 4. LINEAR MIXED MODELS
#------------------------------------------------------------------------------

# Mixed models
lmer_BS       <- lmer(Centroid ~ Env + (1|LineID) + (1|LineID:Env), data = data_final)
lmer_AG       <- lmer(logag ~ Env + (1|LineID) + (1|LineID:Env), data = data_final)
lmer_absAG    <- lmer(logag ~ Env + (1|LineID) + (1|LineID:Env), data = data_filtered)

# Correlation models
lm_cor        <- lm(Centroid ~ logag, data = data_final)
lm_cor_low    <- lm(Centroid ~ logag, data = subset(data_final, Env == "L"))
lm_cor_high   <- lm(Centroid ~ logag, data = subset(data_final, Env == "H"))

lm_cor_low_g  <- lm(Centroid ~ logag, data = subset(cordat, Env == "L"))
lm_cor_high_g <- lm(Centroid ~ logag, data = subset(cordat, Env == "H"))

#------------------------------------------------------------------------------
# 5. MODEL COMPARISONS
#------------------------------------------------------------------------------

lmer_null      <- lmer(logag ~ 0 + (1|LineID), data = data_final)
lmer_first     <- lmer(logag ~ Centroid + (1|LineID), data = data_final)
lmer_BS_AG     <- lmer(logag ~ Env + Centroid + (1|LineID), data = data_final)
lmer_BS_AG_int <- lmer(logag ~ Centroid + Env + (1|LineID) + (1|LineID:Env), data = data_final)

# AIC/BIC
AIC(lmer_null, lmer_first, lmer_BS_AG, lmer_BS_AG_int)
BIC(lmer_null, lmer_first, lmer_BS_AG, lmer_BS_AG_int)

# Likelihood Ratio Tests
anova(lmer_null, lmer_first)
anova(lmer_first, lmer_BS_AG)
anova(lmer_BS_AG, lmer_BS_AG_int)

#------------------------------------------------------------------------------
# 6. RESIDUAL ACCESSORY GLAND SIZE
#------------------------------------------------------------------------------

residual_model <- lm(logag ~ Centroid, data = data_final)
data_final$residuals <- NA
data_final$residuals[as.numeric(rownames(model.frame(residual_model)))] <- residuals(residual_model)

lmer_rag_mixed <- lmer(residuals ~ Env + (1|LineID) + (1|LineID:Env), data = data_final)

#------------------------------------------------------------------------------
# BAYESIAN ANALYSIS: Using Genetic Relatedness Matrix from DGRP
#------------------------------------------------------------------------------

# Download relatedness matrix from DGRP website
url <- "http://dgrp2.gnets.ncsu.edu/data/website/freeze2.common.rel.mat"
local_file <- "freeze2.common.rel.mat"
download.file(url, destfile = local_file, mode = "wb")

# Check file
file.exists(local_file)  # Should return TRUE
lines <- readLines(local_file, n = 10)
print(lines)

# Load matrix
data_mat <- read.table(local_file, header = TRUE, sep = "\t")
rownames(data_mat) <- data_mat[, 1]
data_mat <- data_mat[, -1]
data_mat <- as.matrix(data_mat)

# Clean column and row names
library(dplyr)
colnames(data_mat) <- gsub("\\.line_.*", "", colnames(data_mat))
rownames(data_mat) <- gsub(" line_.*", "", rownames(data_mat))
rownames(data_mat) <- gsub("line_", "dgrp", rownames(data_mat))
colnames(data_mat) <- gsub("line_", "dgrp", colnames(data_mat))

# Match IDs to phenotypic data
unique_ids <- unique(data_final$OriginalID)

# Subset matrix to matching lines
subset_matrix <- data_mat[rownames(data_mat) %in% unique_ids,
                          colnames(data_mat) %in% unique_ids]

# Identify unmatched levels
levels_in_matrix <- unique_ids[!(unique_ids %in% rownames(subset_matrix))]

# Subset data accordingly
data_mcmc <- data_final[!data_final$OriginalID %in% levels_in_matrix, ]
data_mcmc$OriginalID <- as.factor(data_mcmc$OriginalID)

#------------------------------------------------------------------------------
# PREPARE MATRIX FOR MCMCglmm
#------------------------------------------------------------------------------

# Convert to sparse matrix
subset_matrix <- as(subset_matrix, "dgCMatrix")

# Check for positive definiteness
all(eigen(subset_matrix)$values > 0)

# Invert matrix
cor_mat_inverse <- solve(subset_matrix)

#------------------------------------------------------------------------------
# DEFINE PRIORS AND FIT BAYESIAN MODEL
#------------------------------------------------------------------------------

# Empirical prior variance
p.var <- var(data_mcmc$logag, na.rm = TRUE)

prior <- list(
  G = list(G1 = list(V = p.var, nu = 0.002)),  # Random effect for OriginalID
  R = list(V = p.var, nu = 0.002)              # Residual variance
)

# Fit MCMCglmm model
bayesian_model <- MCMCglmm(
  logag ~ Env + Centroid,
  random = ~ OriginalID,
  family = "gaussian",
  rcov = ~ units,
  ginverse = list(OriginalID = cor_mat_inverse),
  data = data_mcmc,
  prior = prior,
  nitt = 100000,
  burnin = 3000,
  thin = 5
)

summary(bayesian_model)

#------------------------------------------------------------------------------
# 8. VISUALIZATION
#------------------------------------------------------------------------------

# Basic plots

data_final |>
  tidyplot(x = Centroid, y = logag, color = Env) |>
  add_data_points() |>
  add_curve_fit(method = "lm")

cordat |>
  tidyplot(x = Centroid, y = logag, color = Env) |>
  add_data_points() |>
  add_curve_fit(method = "lm")

# Long format for traits
data_long <- data_final|>
  pivot_longer(cols = c(logag, Centroid), names_to = "trait", values_to = "value")

# Genotype responses by environment
data_long |>
  mutate(OriginalID = reorder(OriginalID, value, FUN = mean, na.rm = TRUE)) |>
  tidyplot(x = Env, y = value, color = OriginalID) |>
  add_mean_dot(saturation = 2) |>
  add_sd_errorbar() |>
  add_mean_line(color = "red", dodge_width = 0) |>
  split_plot(by = trait)

# Trait distribution by genotype
data_long |>
  mutate(OriginalID = reorder(OriginalID, value, FUN = mean)) |>
  tidyplot(x = OriginalID, y = value) |>
  add_boxplot() |>
  adjust_font(family = "Timesnewroman", face = "plain", fontsize = 2) |>
  adjust_size(width = 125, height = 125) |>
  adjust_x_axis(rotate_labels = TRUE) |>
  split_plot(by = trait)

# Trait distribution by environment
data_long |>
  tidyplot(x = Env, y = value) |>
  add_boxplot() |>
  split_plot(by = trait)

# Residual AG size by environment and genotype
data_final|>
  mutate(OriginalID = reorder(OriginalID, residuals, FUN = mean)) |>
  tidyplot(x = Env, y = residuals, color = OriginalID) |>
  add_mean_dot(saturation = 2) |>
  add_sd_errorbar() |>
  add_mean_line(color = "red", dodge_width = 0)
```
